# Cell strain-derived induced pluripotent stem cell as a genetically controlled approach to investigating aging mechanisms and viral pathogenesis

**DOI:** 10.1101/2021.05.11.443563

**Authors:** Amanda Makha Bifani, Hwee Cheng Tan, Milly M Choy, Eng Eong Ooi

## Abstract

The expansion of the geographic footprint of dengue viruses (DENVs) and their mosquito vectors have affected more than half of the global population, including older adults who appear to show elevated risk of severe dengue. Despite this epidemiological trend, how age and senescence impact virus-host interactions involved in dengue pathogenesis to increase the risk of severe dengue is poorly understood. Herein, we show that conversion of diploid cells with finite lifespan into iPSCs followed by differentiation back into cell strain can be an approach to derive genetically identical cells at different stages of senescence to study virus and aging host interactions. Our findings show that cellular senescence impact the host response to infection and the ensuing outcome. We suggest iPSC-derive cell strains as a potentially useful technical approach to genetically controlled host-virus interaction studies to understand how aging impact viral pathogenesis.

## INTRODUCTION

Dengue is the most common mosquito-borne viral disease globally (1). This acute disease, which when severe can be life-threatening, is caused by four genetically distinct dengue viruses (DENVs) (DENV1,−2,−3 and −4), all of which belong to *Flavivirus* genus. An estimated 390 million infections occur annually (2) and populations throughout the tropics face frequent and recurrent dengue epidemics. More are expected to be affected as the geographic footprint of the *Aedes* mosquitoes that transmit DENV expand from the tropical to the subtropical regions of the world (3).

When frequent and recurrent dengue epidemics first emerged in Southeast Asia after the Second World War, dengue was primarily a paediatric disease (4, 5). Early-life exposure to DENV remains enriched in children in certain parts of the region, leading to immunity by early adulthood (5). However, changes in the urban population demographics as well as vector distribution have led to a shift in the burden of dengue to include adults and even the elderly (6–10). Dengue in older adults present public health challenges as these individuals appear to experience greater morbidity and mortality rates (11). Epidemiological observations have found increased rates of hospital and intensive care admissions (11), length of hospitalisation (12), and risk of severe dengue (12–15). Although age is associated with increased prevalence of co-morbidities, such as cardiovascular diseases and diabetes that also complicate dengue (16, 17), age alone has also been shown to be a risk factor for severe dengue (12). This age-related increased risk of severe disease extends beyond dengue. Vaccination with the live attenuated yellow fever vaccine (YF17D) in those above 60 years of age has, despite the attenuated nature of YF17D, has been associated with severe viscerotropic infection and disease (18, 19).

Despite the increased risk of poor clinical outcome in older adults, how aging affects the pathogenesis of DENV infection and severe adverse events following YF17D vaccination has remained undefined. A major limitation is the lack of suitable *in vitro* tools. Cell lines that are commonly used in virus-host interaction studies are immortal and do not age. Cell strains, or diploid cells with finite lifespan, do age (20). However, most of these cell strains were developed decades ago and are thus mostly close to the end of their finite lifespan. Moreover, global stocks of several of these cell strains are approaching depletion (21). Cell strains at a spectrum of chronological ages are thus not readily available for virus and aging host interaction studies.

Herein, we explored the use of induced pluripotent stem cells (iPSCs) generated from senescent diploid cells, and then differentiated from iPSCs back into senescent cells as a resource for age-dependent viral pathogenesis investigations; conversion of diploid cells to iPSCs serve as a renewable resource for differentiation and passaging into genetically identical cells at different stages of senescence. We show that early passages of differentiated cells display markers of differentiation while later passage cells exhibit cellular senescence. The difference in passage number influences the flavivirus infection phenotype, potentially offering an *in vitro* system to study host immune response to infection in the context of cellular senescence.

## RESULTS

### Senescent cell strains can be reprogrammed into induced pluripotent stem cells

Cell strains, WI-38 and MRC-5 were created as cancer free, virus free cells for vaccine production (20, 22). These diploid cell strains, however, have since proven useful for *in vitro* cell biology and basic virology studies as they are not immortalised (23, 24). We reprogrammed these cell strains into iPSCs using non-modified of Yamanaka factors (Klf4, Oct4, Sx2 and c-Myc) as well as the transcription factors Nanog and Lin28 (25) rather than conventional dedifferentiating techniques such as retro- or lentivirus vectors that alter the host cell genome (Figure 1). Fourteen days post-transfection with the cocktail of reprogramming mRNA, three suspected iPSC colonies were isolated from WI-38 (W1-3) and 6 colonies from MRC-5 cells (M1-6).

**Figure 1.**
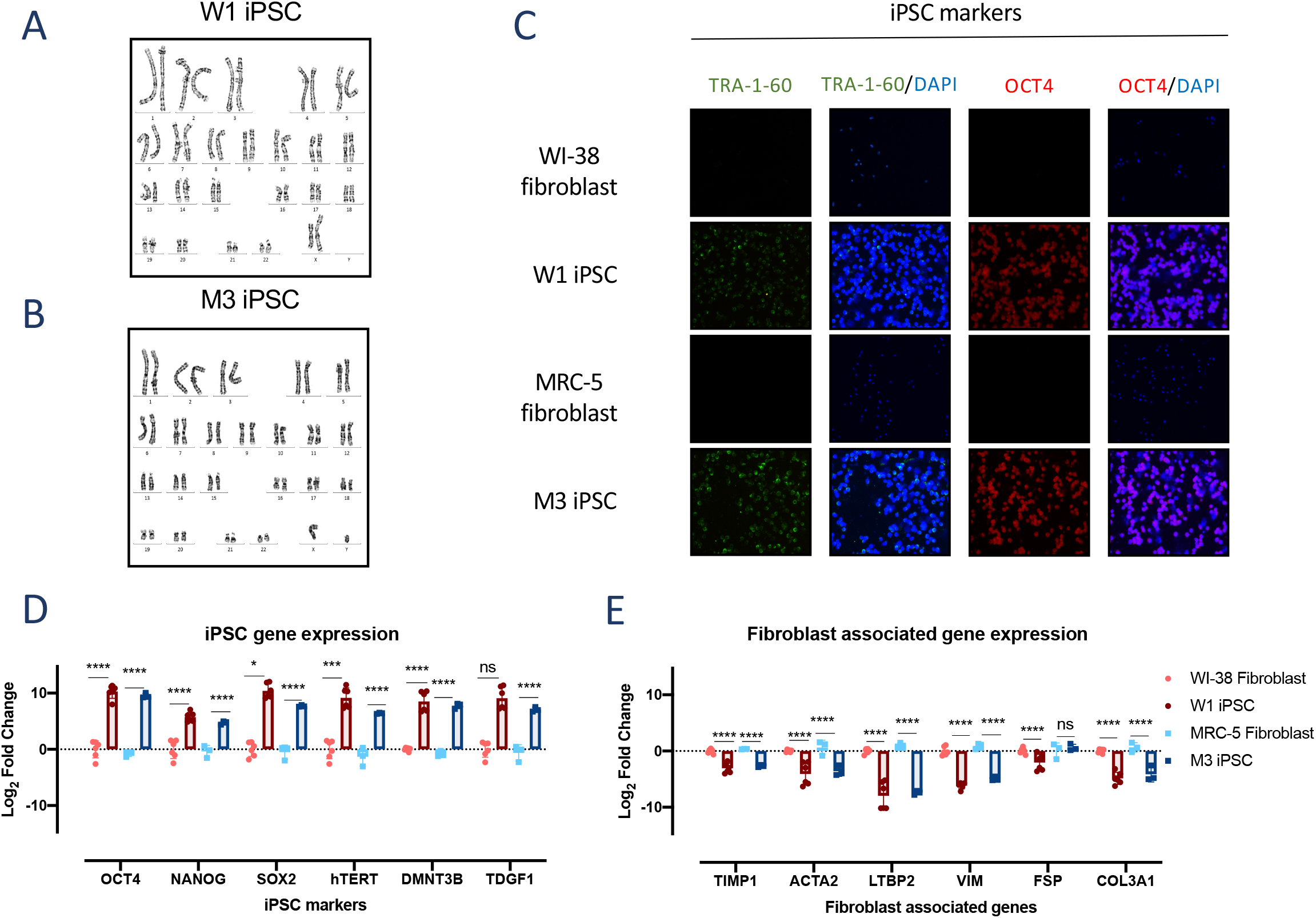
Senescent cell strains WI-38 and MRC-5 can be reprogrammed to iPSCs. (A-B) Karyogram of WI-38 derived iPSC colony W1 (a) and of MRC-5 derived iPSC colony M3 (b) (GTG-banded cells analysed, n=20. Karyograms made, n=5). (C) Immunofluorescence assay employing anti-Tra-1-60 (1/500), anti-oct4 (1/250) and nuclear stain DAPI (1/10000) in parent WI-38 and MRC-5 fibroblasts as well as W1 and M3 iPSCs at 10X magnification. (D-E) Quantitative PCR of fibroblast associated genes and iPSC marker genes in parental cell strains and de-differentiated iPSCs. Statistical analysis was performed using students t-test (each dot represents 1 experiment, n=3 biological replicates/experiment, *p ≤0.05, **p ≤0.01, ***p ≤0.001, ****p ≤0.0001).

To verify that a diploid genome was maintained through reprogramming from senescent fibroblasts to iPSCs, karyotyping was done for all three WI-38 derived colonies and six MRC-5 derived colonies. Colony W1 retained diploid karyotype (Figure 1a) while colonies W2 and W3 exhibited heterogeneous karyotypes (Supplementary Figure S1a). All six colonies isolated from MRC-5 (M1 to M6) preserved their diploid phenotype (Figure 1b; Supplementary Figure S1a).

Colonies W1 and M3, were conveniently selected for further characterisation. We found increased expression of iPSC cell surface marker TRA-1-60 and transcription factor OCT 4 in these colonies (Figure 1c). Reprogramming from fibroblasts to iPSCs was further confirmed at the level of transcription by significantly decreased expression of fibroblast associated genes (*acta2, col3a1, fsp, ltbp2, timp1* and *vim*) and increased expression of iPSC gene markers (*dmnt3b, htert, nanog, oct4, sox2 and tdgf1*) relative to the parental fibroblasts (Figure 1d-e).

A hallmark of cell strains WI-38 and MRC-5 fibroblasts is that they undergo senescence (20) due to lack of expression of human telomerase (hTERT) which prevents telomere shortening. We thus measured hTERT expression in our colonies. Expression of hTERT was upregulated in W1 and M3 iPSCs as compared to their respective parental fibroblasts (Figure 1e).

Stemness was also validated through measuring alkaline phosphatase (AP) activity, which is present in stem but not differentiated cells. Indeed, parental WI-38 fibroblasts stained negative for AP while W1 and M3 iPSCs demonstrated positive pink staining of AP activity (Supplementary Figure S1b).

We further tested if these iPSCs were indeed pluripotent by showing that these cells were capable of differentiation into the three germ layers. The iPSCs were exposed to differentiation media for either endoderm, ectoderm or mesoderm lineages. Lineage differentiation was confirmed based on increased expression of lineage specific markers (endoderm: *sox17, gata6, foxa2;* mesoderm: *ncam1, hand1, msx1;* ectoderm: *otx2, pax6, lhx2*) (Supplementary figure S1c) matched by decreased expression of iPSC markers (*nanog, sox2, oct4*) (Supplementary figure S1d). Taken collectively, these multiple lines of evidence suggested that WI-38 and MRC-5 were successfully reprogrammed to W1 and M3 iPSCs.

### IPSC-derived differentiated cells undergo senescence

To derive differentiated cells from iPSCs, we added a chemically defined media to the iPSCs and passaged the cells four times (Figure 2a). Stem cell morphology was lost upon differentiation from iPSC to a differentiated cell (Figure 2b). Furthermore, there was a significant decrease in iPSC marker genes upon differentiation in both W1 and M3 differentiated cells at every passage post differentiation (Figure 2c-d). However, after four passages, the cells could no longer be maintained in culture and perished.

**Figure 2.**
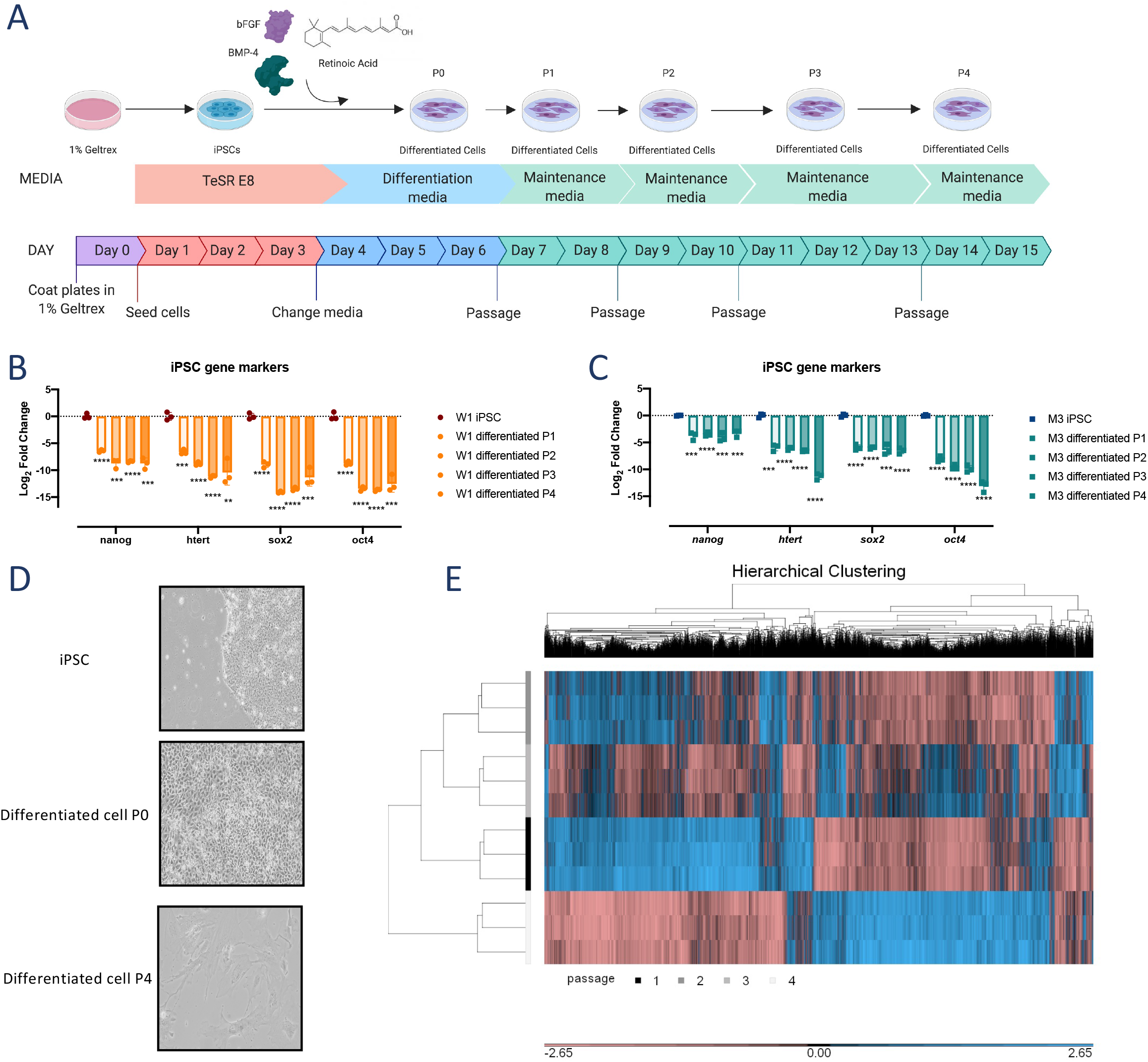
IPSCs can be differentiated back into cell strains. (A) Schematic depicting the protocol for define differentiation of iPSCs. Image made in BioRender. (B-C) Quantitative PCR of iPSC marker genes in W1 (B) and M3 (C) iPSCs and the differentiated cells derived from these respective iPSCs, at P1-P4. (D) Bright field images of W1 cells prior to differentiation, during differentiation and following differentiation at passage 1 and 4 at 10X magnification. (E) Hierarchical clustering of microarray gene expression from passage 1 to 4 of W1-derived differentiated cells. *p ≤0.05, **p ≤0.01, ***p ≤0.001, ****p ≤0.0001.

To gain insights into the differences between passage 1 (P1) to P4 differentiated cells, we analysed whole genome expression of these cells using microarray. Hierarchical clustering of differentially expressed genes from P1 through P4 of W1 differentiated cells revealed distinct patterns at P1 and P4 (Figure 2e). Gene expression patterns at P2 and P3 were intermediate to those of P1 and P4 (Figure 2e).

Pathway analysis of genes differentially expressed at P2, P3 and P4 compared to P1 defined the differences between the cells at different passages. Few genes were significantly (adjusted p-value < 0.05) up (n = 14) or down (n=85) regulated from P1 to P2. The top hits of upregulated genes were related to cell differentiation processes (Figure 3a), whereas the downregulated genes did not significantly associate with a specific pathway (Supplementary figure 2a). The comparison between P1 against P3 and P4 cells yielded interesting findings. Senescence and autophagy in cancer cells emerged as the second top pathway in both analyses (Figure 3a). Other aging-related pathways that were highlighted in our analysis were complement and coagulation cascades, hypothesized pathways in cardiovascular disease and genotoxicity. Interestingly, acute inflammation was also upregulated at P4 (Figure 3a), which is supportive of the notion of inflammaging - an increase in basal inflammation in elderly individuals (26). These findings were matched with significant downregulation of pathways associated with DNA replication and the cell cycle (Supplementary figure 2c). Differential expression of genes associated with senescence (*serpine1*) and acute inflammation (*icam1*) were validated by qPCR of W1 and M3 differentiated cells at P1 and P4 (Supplementary figure 2d).

**Figure 3.**
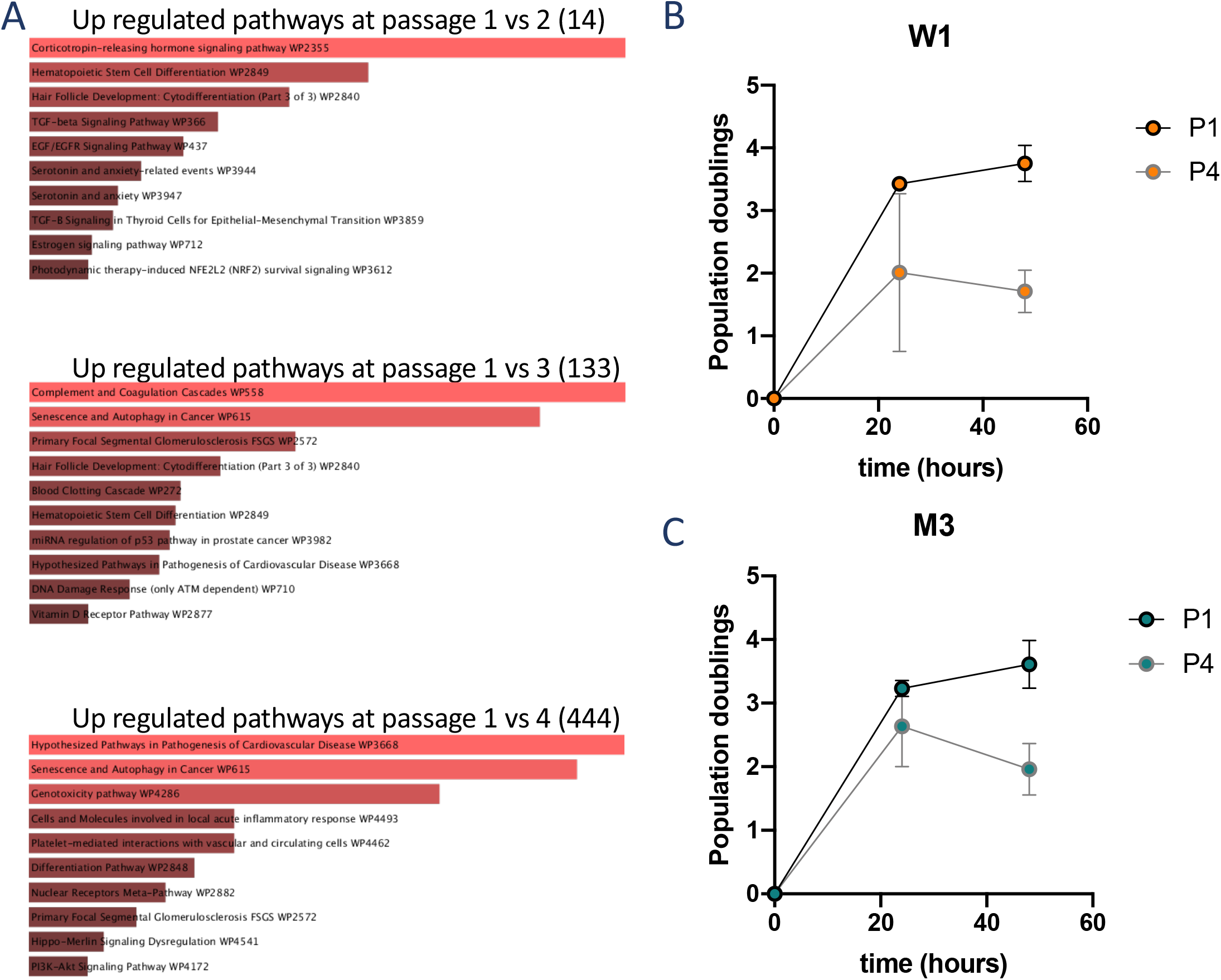
IPSC-derived differentiated cells undergo senescence. (A) Pathway analysis of upregulated genes in P1 vs either P2, P3 or P4 of W1 differentiated cells using the WikiPathways 2019 Human database. Pathways are shown in order of significance, with the most significantly upregulated pathway at the top. (B-C) Population doubling time of W1 and M3 differentiated cells at P1 and P4 over 48 hours of observation.

The senescent phenotype was phenotypically validated by measuring population doubling times at passage one and four in differentiated cells. Aging cells have previously been shown to have slower population doubling times. Both W1 and M3-derived differentiated cell demonstrated slower population doubling time at later passages (Figure 3b-c).

### Differentiated cells but not iPSCs are susceptible to DENV infection

Stem cells have recently been shown to be less susceptible to viral infection through intrinsic expression of interferon stimulated genes (ISGs) independent of interferonβ (IFNβ) activity (27). We thus measured a selection of these previously identified ISGs by qPCR in our iPSCs and compared to their parental fibroblasts to ensure that our iPSCs showed the same ISG expression profile as those previously reported (27). We found that the ISGs - *alyfref, eif3l, pabpc4, ptma* and *ybx3* - were intrinsically expressed and were significantly upregulated in the iPSCs compared to their parental fibroblasts (Figure 4a).

**Figure 4.**
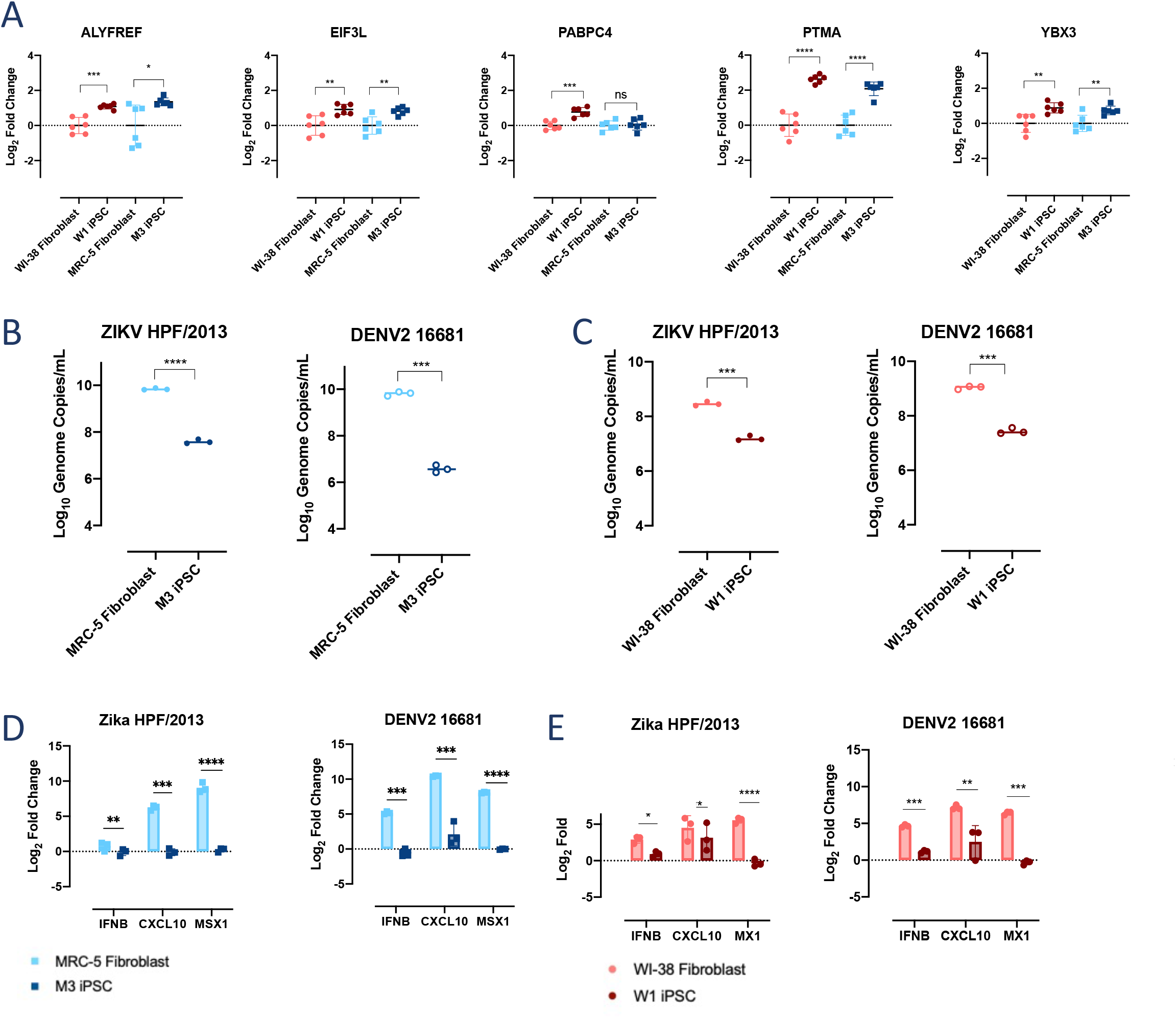
IPSCs are resistant to flaviviral infection through an IFNβ independent mechanism. (A) Quantitative PCR of ALYFREF, EIF3L, PABPC4, PTMA and YBX3 (stem cell interferon independent ISGs) in W1 and M3 iPSCs. Their respective parental cell strains were included as controls. (B-C) RT-PCR of genome copies of DENV2 16681 and ZIKV HPF/2013 from the supernatant of infected cell strains and iPSCs at 72 hours post infection (hpi) in M3 and W1 iPSCs. (D-E) Quantitative PCR of interferon and interferon stimulated genes in iPSCs and parental cell strains MRC-5 (D) and WI-38(E) infected with ZIKV (HPF/2013) or DENV2 (16681) relative to the respective uninfected control cell.

Along with the intrinsically expressed ISGs, our iPSCs were less susceptible to DENV and ZIKV infection. Inoculation of wild type DENV2 (16681 strain) and Zika virus (ZIKV) (HPF/2013 strain) onto iPSCs resulted in significantly reduced viral genome copies at 72 hours post infection (hpi) compared to infection in their respective parental fibroblasts (Figure 4b-c). Similar differences were observed when infection was assayed for infectious viral progenies (Supplementary figure 3a). Furthermore, there was a significantly attenuated type I interferon response during infection with both viruses in both iPSCs relative to the original fibroblasts (Figure 4d-e; Supplementary figure 3b-c), consistent with previous observations (27).

We next asked if cell susceptibility to viral infection could be restored in iPSC-derived differentiated cells. W1 differentiated cells were infected with DENV2 16681, while M3 differentiated cells were infected at passages 1 and 4. We found infectious DENV2 particles in the supernatant of W1 and M3 differentiated cells at 48 hpi (Figure 5a); the plaque titres were, however, not significantly different. DENV2 genomic RNA in the supernatant of infected W1 cells showed an increasing trend with increasing number of passages although this difference was also not statistically significant (Figure 5b). No difference was seen in DENV2 16681 RNA levels in P1 compared to P4 M3 cells (Figure 5b). These findings thus indicate that susceptibility to DENV infection was restored in iPSC-derived differentiated cells.

**Figure 5.**
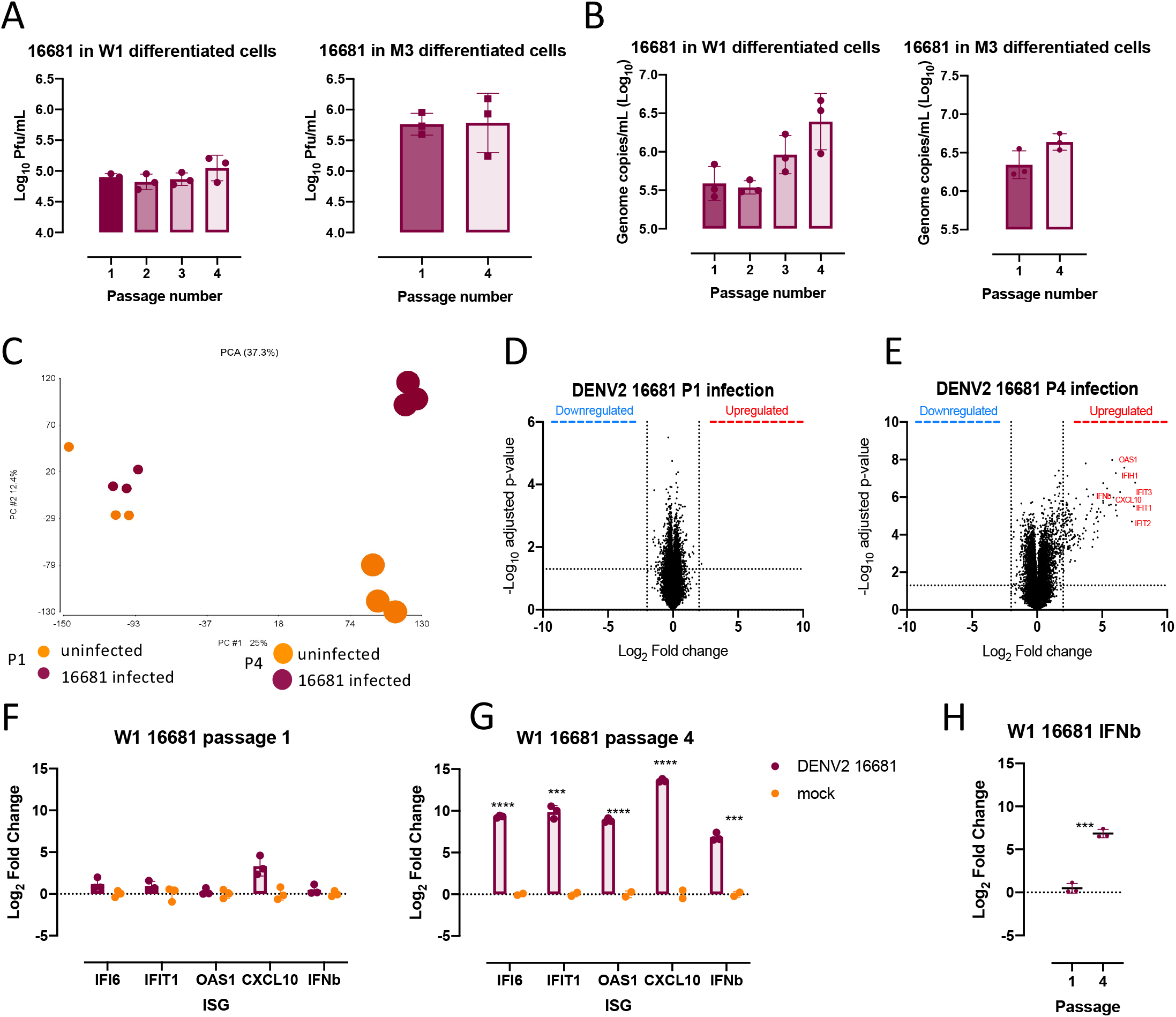
Susceptibility to DENV infection is restored in iPSC-derived differentiated cells. (A) DENV2 16681 progenies produced at 48hpi in P1-P4 W1-derived differentiated cells and M3 differentiated cells at P1 and P4. (B) Genomic RNA recovered from the supernatant of DENV2 16681 infections at 48hpi in W1- and M3-derived differentiated cells. (C) Principal component analysis (PCA) of gene expression data from W1 differentiated cells infected with DENV2 or mock infected at P1 and P4. P1 cells are shown in smaller dots whereas P4 are larger. (D-E) Volcano plot of gene expression changes in DENV2 16681 infected W1-derived differentiated cells compared to mock infection at P1 and P4. Values above the horizontal line are statistically significant. Vertical lines depict the Log_2_ cut off assigned to genes that are differentially expressed. (F-G) Quantitative PCR of iPSC antiviral genes identified in the microarray analysis (IFIT1, IFI6, IFNβ OAS1, CXCL10) during infection with DENV2 16681 at P1 and P4 in infected and uninfected cells. (H) IFNβ expression in infected W1 differentiated cells at P1 vs P4.

### Aging cell related differences in host response to DENV infection

Given the differences in baseline gene expression upon passage of these iPSC-derived differentiated cells (Figure 3a), we next explored if the host response to DENV2 16681 infection was different despite producing similar levels of viral progenies. Gene expression of both infected and mock infected P1 cells clustered together. In contrast, infected P4 cells clustered separately from their mock infected controls (Figure 5c). Only one single gene of unknown function was significantly upregulated in response to DENV2 16681 infection in P1 cells (Figure 5d). This finding is interesting as P1 cells partially resembled CD33+ myeloid cells and would be expected to be more prone to pro-inflammatory response, if the host response to infection was influenced more by cell type rather than age. In contrast, 354 genes were found to be differentially expressed between DENV2 16681 infected and mock infected P4 cells (Figure 5e). Many of these genes showed a large fold change, especially those that belong to the canonical IFNβ antiviral response pathway (Figure 5e). The differentially expressed IFNβ and related genes identified in the microarray analysis were validated through qPCR (Figure 5f-g). Indeed, IFNβ expression was significantly lower in P1 compared to P4 DENV2 infected cells (Figure 5h). The increase in antiviral response at later passages was also observed by qPCR in M3 differentiated cells (Supplementary figure 4).

To determine if the response of the differentiated cells to infection was generic or specific to DENV2 16681, we examined infection outcome in our iPSC-derived differentiated cells using the attenuated DENV2 PDK53 and YF17D. DENV2 PDK53 was derived from its wild-type 16681 parent through *in vitro* serial passaging. Its genome is thus composed of the 16681 genome but with 5 amino acid and 1 nucleotide substitutions, the latter in the 5’ untranslated region of the genome. Despite these small number of genomic differences, infection with PDK53 produced reduced levels of infectious progenies in higher passaged cells (Figure 6a). Furthermore, unlike 16681 infected P1 cells that showed minimal transcriptional level changes, PDK53 infection induced IFNβ expression that was greater than 6 Log_2_fold change compared to mock infected P1 and P4 cells (Figure 6b). Similar trends were also observed with the expression of ISGs (Figure 6c-d, Supplementary figure 5a-b).

**Figure 6.**
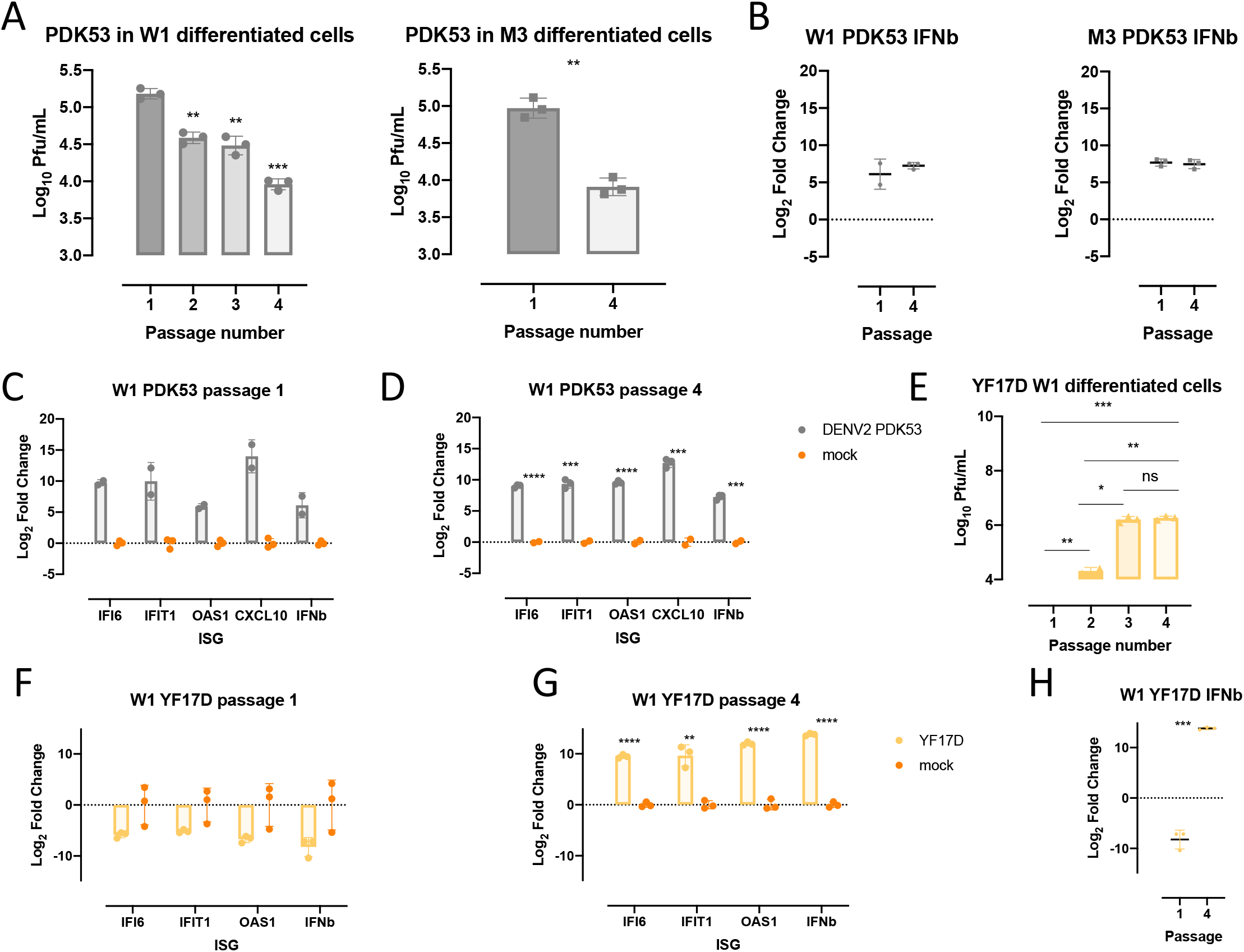
Infection outcome in iPSC-derived differentiated cells is virus-specific. (A) Infectious DENV2 PDK53 particles 48hpi in W1 differentiated cells at passage 1 through 4 and M3 differentiated cells at passage 1 and 4. (B) IFNβ expression in P1 and P4 W1- and M3-derived differentiated cells infected with PDK53. (C-D) IFIT1, IFI6, IFNβ, OAS1, CXCL10 gene expression upon DENV2 PDK53 compared to mock infection in P1 (C) and P4 (D) M3-derived differentiated cells. (E) YF17D infectious viral progeny recovered at 48 hpi in P1-P4 W1-derived differentiated cells. (F-G) W1-derived differentiated cells IFIT1, IFI6, IFNβ, OAS1, CXCL10 gene expression at 48 hours post YF17D infection. (H) YF17D infected W1-differentiated cell’s IFNβ expression at P1 vs P4.

Infection with the YF17D produced the opposite trend compared to PDK53. YF17D infection showed increased virus replication with increasing passage number of differentiated cells (Figure 6e; Supplementary figure 5c) despite increased expression of IFNβ and ISGs (Figure 5f-h; Supplementary figure 5d-f). This observation is particularly interesting as YF17D vaccination is known to be associated with increased risk of viscerotropic infection and disease in older adults (28). Collectively, our findings suggest that the host response to infection is not generic in our iPSC-derived differentiated cells but are rather age-dependent and could potentially inform on age-related host response to flaviviral infection.

## DISCUSSION

Dengue in older adults have shown worse clinical outcome compared to their younger counterparts (11). DENV infection in elderly patients often presents atypically and complicates clinical diagnosis (8, 15, 29). They are also more likely to have had prior exposure to DENV and are thus at greater risk of antibody-dependent enhancement upon secondary infection with a heterologous DENV (30) (31). The prevalence of co-morbidities, such as diabetes and hypertension, increase with age and several of these have been linked with increased risk of severe dengue (16, 17, 32). However, there could also exist age-related host factors, including immunosenescence, the decline of the immune system with age, or inflammaging, an increase in basal inflammation with age, that may elevate the risk of older individuals to severe dengue (33). Thus, despite the known poorer clinical outcome of dengue in older adults, the pathogenic underpinnings of DENV infection in aged cells have remained undefined.

A major limiting step to understanding age-related differences in host-virus interactions is the lack of suitable *in vitro* tools to simulate senescence. Commonly used cell lines are immortal and thus poorly reflect the processes of aging. Cell strains, such as WI-38 and MRC-5 undergo senescence (20, 24). Age associated changes in gene expression may influence outcome of DENV infection. Indeed senescent monocytes have been shown to have increased DENV susceptibility via increased expression of receptor DC-SIGN (34). Moreover, cell strains also have the advantage of having diploid genomes that could be more accurate than cell lines in reflecting the transcriptional responses that happens in dengue patients. Unfortunately, due to their limited lifespan, global stocks of these diploid cell strains are limited. WI-38 diploid fibroblasts have been used to near exhaustion since their isolation. Thus, remaining supplies are constrained to high passaged senescent cells (21, 24). Without ready access to paired low and high passaged cells, WI-38 and MRC-5 have limited potential as tools to dissect age-related host-virus interactions that underpin pathogenesis. Our work thus overcomes this limitation through the derivation of iPSCs from WI-38 and MRC-5. The reprogrammed iPSCs could be differentiated and passaged for infection experiments.

We have used a chemically defined media to differentiate the iPSCs into an adherent cell monolayer, that was followed through serial passaging. We have used this relatively straightforward approach as a proof-of-concept demonstration of WI-38 and MRC-5 derived iPSCs and iPSC-derived differentiated cells as *in vitro* models to study age-related effects on viral infection. Indeed, our transcriptional analysis showed that the baseline expression of aging-related genes were increased upon passaging of W1 and M3 cells. To our knowledge, such an approach to derive isogenic cells of different replicative ages for infection studies has not been previously attempted. Future studies could make use of better defined differentiation protocols using well established kits. Alternatively, iPSCs could also be differentiated through the use of transcriptions factors computationally predicted by mogrify (35) or epimogrify (36). Furthermore, as the process of differentiation does not occur uniformly throughout a culture, single-cell sequencing would enable us to compare homogenous cell types within a heterogeneous population at various replicative ages.

We found interesting differences in the virologic outcome and host response to infection in both W1- and M3-derived cells. DENV2 16681 infection produced no difference in the amount of progeny virus across the different passages iPSC-derived differentiated cells. Conversely, infection with its attenuated derivative, DENV2 PDK53, produced significantly reduced viral progenies with increasing number of passages. This finding is interesting as we have previously found DENV2 PDK53 infection to be restricted by the innate immune response, which may also explain its attenuated phenotype (37, 38). Despite the production of comparable levels of DENV2 16681, infection with this virus in P1 to P4 iPSC-derived differentiated cells produced vastly different responses in gene expression. P4 cells produced more genes with greater fold change in infected compared to uninfected cells than P1 cells. Many of these genes, such as *il1a* or *serpine1*, are in the innate immune and pro-inflammatory pathways. These findings suggest that for any given viral load, older adults may respond differently compared to the younger dengue patients with a more pronounced inflammatory response and hence explain their increased risk in severe dengue.

The possibility that this *in vitro* approach could reflect, at least partially, age-related clinical outcome of infection is further supported by our observations on YF17D infection. Vaccination of individuals over 60 years of age with YF17D has been linked with severe adverse events (18) and viscerotropic disease (28), the latter possibly explained by increased burden of live attenuated YF17D infection. YF17D infection in our passaged cells produced higher levels of infectious particles with increasing number of passages. The increase in YF17D progenies occurred despite increased IFNβ expression at later passages. This observation suggests that other YF17D-host interactions may underpin infection outcome in aged cells and hence alter their risk of severe adverse events following YF17D vaccination.

In conclusion, our findings suggest the feasibility of using iPSC-derivatives of WI-38 and MRC-5 cell strains as a resource to elucidate how aging impact host-virus interactions that underpin dengue and other flaviviral pathogenesis.

## MATERIALS AND METHODS

### Cells and culture conditions

Human diploid fibroblast WI-38 (female) and MRC-5 (male) cell strains were maintained in fibroblasts growth media (Minimum Essential Media, 10% FCS, 1% GlutaMAX, 1% penicillin/streptomycin) at 37°C, 20% O_2_, 5% CO_2_. Cell strains were passaged with TrypLE™ Expression Enzyme. BHK21 cells used for plaque assay were grown in RMPI Medium 1640 (Gibco), 2% FCS and 1% penicillin/streptomycin at 37°C, 20% O_2_, 5% CO_2_.

All stem cells were cultivated on 1% Geltrex™ coated cell culture ware in mTeSR™1 or TeSR™-E8™ media for maintenance. Stem cells were passaged according to manufacturers’ recommendations with ACCUTASE™ or ReLeSR™ where appropriate. Briefly, spent media was removed from the stem cells and ReLeSR™ was added and incubated at room temperature for one minute. ReLeSR™ was then removed, cells were incubated at 37°C, 20% O_2_, 5% CO_2_ for 6:30min, fresh media was added gently, and cells were re-suspended by tapping the plate for 1min. ACCUTASE™ was added to cells and incubated for 7min at 37°C, 20% O_2_, 5% CO_2_. Cells were resuspended, transferred to a conical tube and spun at 250 *x* g for 5min at room temperature. The ACCUTASE™ was then removed and cells were re-suspended in desired media with 10μM ROCK inhibitor Y-27632.

### Virus stocks

Dengue strains (DENV2 16681 and DENV2 PDK53) were gifted by Dr Claire Huang (Centre for disease control and prevention, USA). Clinical isolate Zika virus HPF13/2013 (KJ776791) was acquired by the European Virus Archive. Yellow fever YF17D was isolated from a vial of Stamaril^®^ live attenuated vaccine. All flavivirus stocks were maintained in insect C6/36 cells at 30°C.

### Reprogramming diploid fibroblasts to iPSCs

Human diploid fibroblast cell strains WI-38 and MRC-5 were reprogrammed to iPSCs using the StemRNA^™^-NM reprogramming kit according to manufacturers’ instructions for adult and neonatal human fibroblasts. Briefly, cell strains were seeded on 6 well plates coated with 1% Geltrex™ LDEV-Free Reduced Growth Factor Basement Membrane Matrix at a density of 2 x 10^5^ cells per well in fibroblast expansion media (Advanced DMEM, 10% FCS, 1% Glutamax) and incubated at 20% O_2_, 37°C overnight. Subsequently, spent media was replaced with NutriStem Media and incubated in at 37°C for 6 hours prior to introduction of the NM-RNA reprogramming cocktail with Lipofectamine^®^RNAiMAX^™^ Transfection Reagent in Opti-MEM^®^ Reduced Serum Medium. Fresh Nutristem Media and NM-RNA reprogramming cocktail was refreshed daily for the course of four days. Cells were subsequently maintained in Nurtistem media until iPSC colonies could be identified (~14 days). Contrary to manufacturer’s instructions, reprogramming was carried out at 20% O_2_ rather than ≤5% O_2_.

Potential colonies of iPSCs were manually isolated using a micropipette with a 20μl tip and transferred to a fresh 1% Geltrex™ coated 6 well plate containing mTeSR^™^1 maintenance media for propagation according to manufacturers’ instructions.

### Gene expression quantification

Relative changes in gene expression of lineage specific markers were measured by qPCR. Briefly, cellular RNA was isolated following the RNeasy Mini Kit, and converted to cDNA via qScript standard protocol. QPCR were performed using LightCycler^®^ 480 SYBR Green I Master under the conditions on LightCycler^®^ 480 II using LightCycler^®^ 480 software (v.1.5). Gene expression primers can be found in supplementary table 1.

### Immunofluorescent assay

Primary antibodies against iPSC markers TRA-1-60 (ab16288, Abcam^®^, 1/500) and OCT4 (ab181557, Abcam^®^, 1/250) were used for immunofluorescence assays to determine the expression of stem cell proteins in iPSCs or fibroblasts. Spent media was removed, cells were washed once with PBS and subsequently dislodged using ACCUTASE™ according to manufacturers’ instructions. The cells were spun down at 250 *×* g for 5min at room temperature and the supernatant was decanted. Pelleted cells were re-suspended in 250μL PBS. Two microliters of re-suspended cells were aliquoted on to 30 well microscope slides (TEKDON incorporated, Slide ID:30-30), allowed to air dry and fixed in acetone for 10min at room temperature. Slides were washed in a 50mL conical tube containing PBS for 5min at room temperature three times before primary antibody was applied and incubated for 1-2 hours at 37°C. The slides were washed three times with PBS at room temperature for 5 minutes. Anti-mouse or anti-rabbit secondary antibody was applied where appropriate and incubated for 30 minutes at 37°C. Slides were rinsed with PBS at room temperature for 5 minutes three times. The SlowFade™ Antifade Kit was used as a mounting medium as well as to stain the cellular DNA with DAPI. Slides were visualised on a Nikon Eclipse 80*i* microscope with Nikon Intensilight C-HGFI at 10X magnification and imaged with Nikon Digital Sight camera using NIS Elements Imaging Software (v.3.22.15).

### Trilineage differentiation of iPSCs

Pluripotency was confirmed using STEMdiff^™^ Trilineage Differentiation Kit according to manufacturers’ instructions. Stem cells were harvested using ACCUTASE™ and seeded on 24 well plates coated with 1% Geltrex at 1 x 10^5^ cells per well for mesoderm differentiation and 1 x 10^5^ cells per well for ectoderm and endoderm differentiation in their respective differentiation medium. Differentiation to endoderm, ectoderm and mesoderm was assessed by qPCR as described earlier using lineage specific primers (Supplementary table 1). Differentiated cells gene expression was assessed against undifferentiated iPSC control.

### Stem cell differentiation

The spent medium of W1 iPSCs was removed and the cells were treated with ACCUTASE™ and incubated for 7min at 37°C, 20% O_2_, 5% CO_2_. Cells were dislodged, transferred to a 15mL conical tube, spun at 250 *×* g for 5 min at room temperature, the supernatant was decanted and pelleted cells were re-suspended in 3mL TeSR™-E8™ in the presence of 10μM Y-27632. Cells were later seeded on 6 wells plates coated with 1% Geltrex™ at a density of 2 x 10^5^ cells/well in TeSR™-E8™ supplemented with Y-27632 and kept at 37°C, 5% CO_2_. Two days later, spent medium was removed and replaced with differentiation medium (DMEM/F12, 1% ITS, 0.001nM isoprenaline, 100ng/mL BMP-4, 20ng/mL bFGF and 0.1μM Retinoic acid) and incubated at 37°C, 20% O_2_, 5% CO_2_ for four days. Cells were then imaged and maintained in culture or the RNA was extracted for qPCR to assess iPSC markers as previously described. Differentiated cells were maintained in DMEM supplemented with 1% Penicillin/streptomycin, 10% Foetal calf serum, 1mM L-glutamine at 37°C, 5% CO_2_ thereafter.

### Alkaline Phosphatase Activity

Undifferentiated cells were characterized by upregulated alkaline phosphatase activity compared to terminally differentiated cells. Human diploid fibroblasts and iPSCs were seeded in triplicates at 1 x 10^5^ cells/well of fibroblasts or 200 clumps/well for the iPSCs. Alkaline phosphatase activity was measured using the StemAb Alkaline Phosphatase Staining Kit II in accordance with manufactures’ guidelines. Cells were imaged on an Olympus DP71 with Olympus TH4-200 camera and recorded with CellSens imaging software.

### Viral infections

Infections were performed on senescent cells strains, iPSCs and differentiated with dengue 2 wild type strain 16681, dengue 2 vaccine strain PDK53, Zika wild type stain HPF/2013 or yellow fever vaccine strain 17D. Fibroblasts were seeded in a 24 well plate and kept at 37°C, 20% O_2_, 5% CO_2_ overnight. Spent medium was removed, cells were counted and infected at an MOI of 1 for 1hour at 37°C, 5% CO_2_ in MRC-5 and WI-38 fibroblasts as well as their respective iPSCs. Differentiated cells were infected with virus at an MOI of 0.1 for 1 hour in 37°C, 5% CO_2_. Virus inoculum was removed and replaced with fresh media. Fibroblasts and differentiated cells were incubated at 37°C, 5% CO_2_ for 48 hours while iPSCs were incubated for 72 hours. The supernatant was harvested for plaque assay and qRT-PCR of the viral genome. Cells were collected for RNA extraction followed by qRT-PCR.

### Infection quantification

Infectious particle quantification was determined via plaque assay. Briefly, BHK21 cells were seeded at 2 x 10^5^ cells/well in a 24 well plate in RMPI medium supplemented with 2% FCS and 1% penicillin/streptomycin. Once confluency was reached, the BHK21 cells were infected with a 10-fold serial dilution of 100μL of virus and incubated at 37°C, 20% O_2_, 5% CO_2_ for 1 hour, with plate agitation at 15 minute intervals. Subsequently, viral inoculum was removed, 0.8% carboxymethyl cellulose (CMC) in RPMI medium supplemented with 3% FCS and penicillin/streptomycin was added and plates were incubated at 37°C, 20% O_2_, 5% CO_2_ for six days. Cells were then fixed in 20% formalin for at least 30 minutes before rinsing with water. Plates treated with 1% crystal violet (Sigma-Aldrich), washed and air-dried to count plaques.

Viral genome copy number was quantified by qRT-PCR using CDC primers and probes as previously described for dengue virus (39) and zika virus (40). Briefly, virus genomic RNA was extracted from the supernatant using QIAamp Viral RNA Mini Kit according to manufacturer’s instructions. Purified RNA was then quantified by qPCR following the qScript One-Step RT-PCR Kit protocol using the CDC specified primers and probes on the LightCycler^®^ 480 II.

### Microarray analysis

RNA was extracted from infected and uninfected differentiated cells at 48 hours post infection using the RNeasy Micro Kit. Micrroarray analysis was done using the GeneChip™ Human Gene 2.0 ST array. Analysis was carried out using the Partek^®^ gene expression suit. A list of differentially expressed genes were selected based on a FDR adjusted p-value of <0.05 and a gene expression difference of at least Log_2_fold-change of 2. Hierarchical clustering was carried out on the genes lists generated. Pathway analysis was done on Enrichr using WikiPathways.

### Population doubling of differentiated cells

Cells were seeded at 5 x 10^4^ cells/well in a 24 well plate. At 24 hours and 48 hours, cells were washed with PBS and treated with trypsin. The dislodged cells were counted in triplicate. Population doubling time was calculated from the number of cells at 0, 24 and 48 hours post seeding.

### Quantification and statistical analysis

All statistical analyses were performed using GraphPad Prism (v.8.2.1). Student t-test was used and *p*-value of ≤ 0.05 was considered significant (ns > 0.05, \**p ≤* 0.05, \*\**p ≤* 0.01, \*\*\**p ≤* 0.001, \*\*\*\**p ≤* 0.0001). Statistics depicted on graphs show the mean and standard deviation about the mean unless stated otherwise. All data points shown are biological replicates, unless otherwise stated in the figure legend.

## ACKNOWLEDGEMENTS

The authors would like to thank Claire Huang from the Centers for Diseases Control and Prevention gifting the DENV strains used in this paper. AMB is supported by a graduate studentship from Duke-NUS Medical School. EEO receives salary support from the National Medical Research Council of Singapore, through the Clinician-Scientist (Senior Investigator) Award.

## Supplementary Figures Legends

**Supplementary figure 1.**
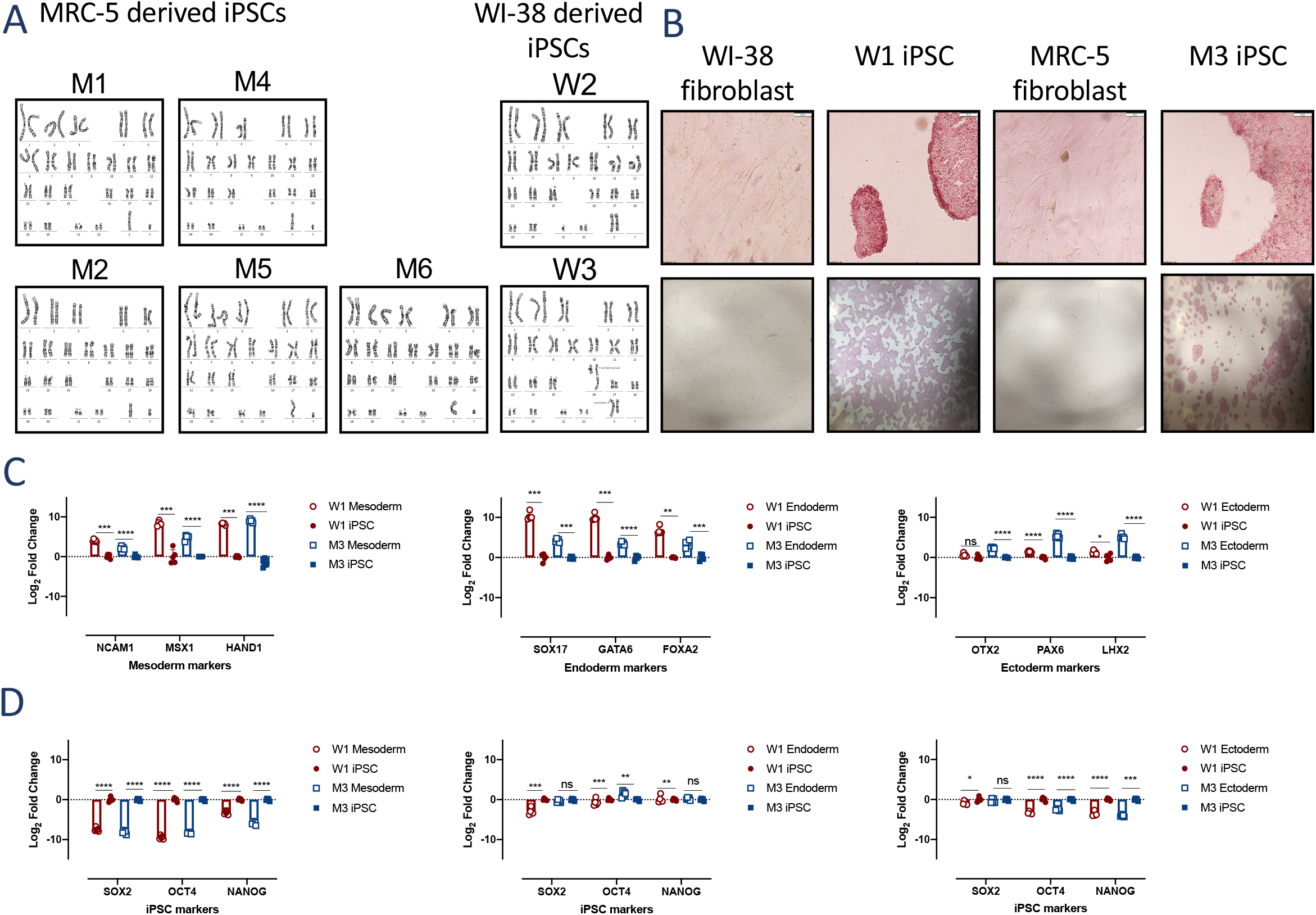
Senescent fibroblast reprogrammed into iPSCs. **(A)** MRC-5 derived iPSC colonies M1, M2, M4, M5 and M6 karyograms depicting normal male karyotype. Aberrant karyotype of WI-38 derived iPSC colony W2 and W3 with anomalies in translocation or chromosome copy number indicated on the karyogram. (GTG-banded cells analyzed, n=20. Karyograms made, n=5). **(B)** Alkaline phosphatase staining in WI-38 and W1 iPSC as well as MRC-5 and M3 iPSC. Cells with elevated phosphatase activity stained red/pink. **(C-D)** Quantitative PCR analysis of tri-lineage differentiation of W1 and M3 iPSCs into mesoderm, endoderm and ectoderm using appropriate lineage specific markers (C) and iPSC markers (D). Student t-test (n = 3) was used for statistical analysis *\*p ≤*0.05, \*\**p ≤*0.01, \*\*\**p ≤* 0.001, \*\*\*\**p ≤* 0.0001.

**Supplementary figure 2.**
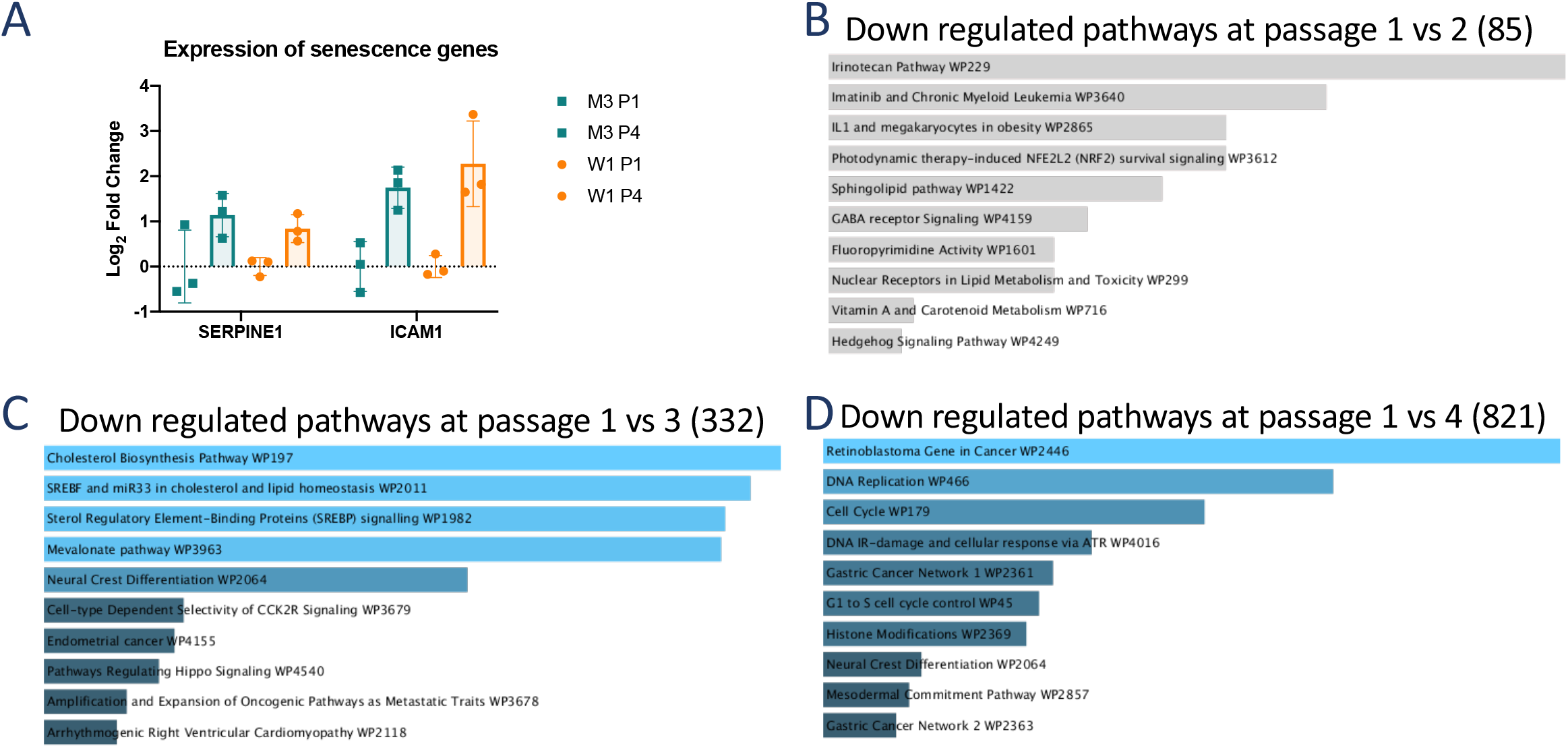
Differentiated cells exhibit hallmarks of aging. **(A)** Quantitative PCR of senescence associated genes in W1- and M3-derived differentiated cells at P1 and P4 (each dot represents 1 replicate). **(B-D)** Pathway analysis of downregulated genes in P2 (B), P3 (C) and P4(D) of W1 differentiated cells compared to P1 using the WikiPathways 2019 Human database. Significantly enriched downregulated pathways are coloured blue with the most significant ones shown at the top. Pathways that did not meet statistical significance of *p < 0.05* are shown in gray.

**Supplementary figure 3.**
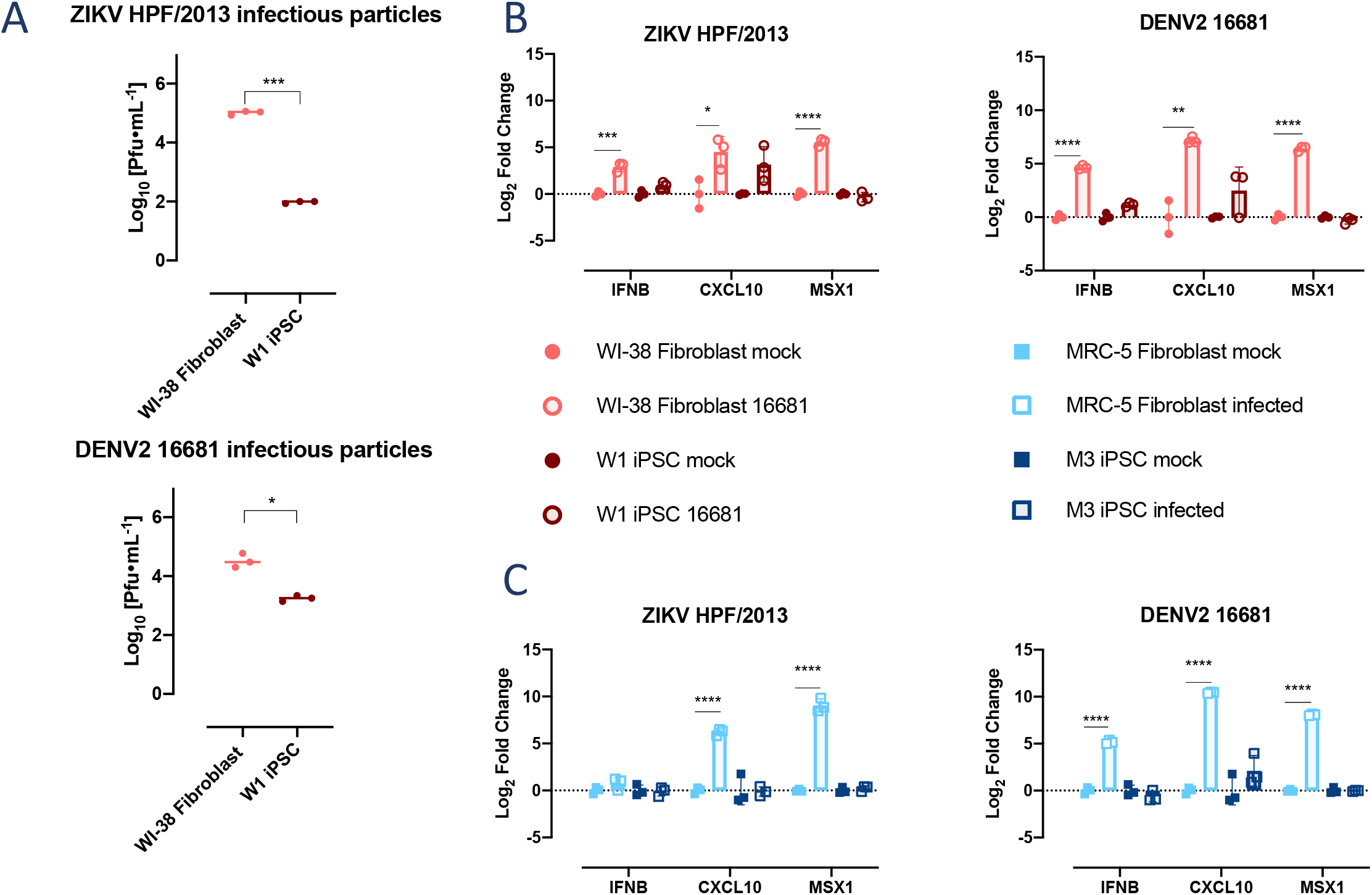
Flaviviral infection and IFNβ response in cells strains and cell strain derived iPSCs at 72 hpi. Cell strains produce more infectious particles and a larger type I IFN response than their iPSC derivatives. **(A)** Zika HPF/2013 and DENV2 16681 infectious particles produced in WI-38 and W1 iPSCs at 72 hpi. **(B-C)** Quantitative PCR of IFN and ISGs in W1 (B) and M3 (C) iPSCs as well as their respective parental cell strains with or without ZIKV and DENV infection. Student t-test (n = 3) was used for statistical analysis \**p ≤* 0.05, \*\**p ≤* 0.01, \*\*\**p ≤* 0.001, \*\*\*\**p ≤* 0.0001

**Supplementary figure 4.**
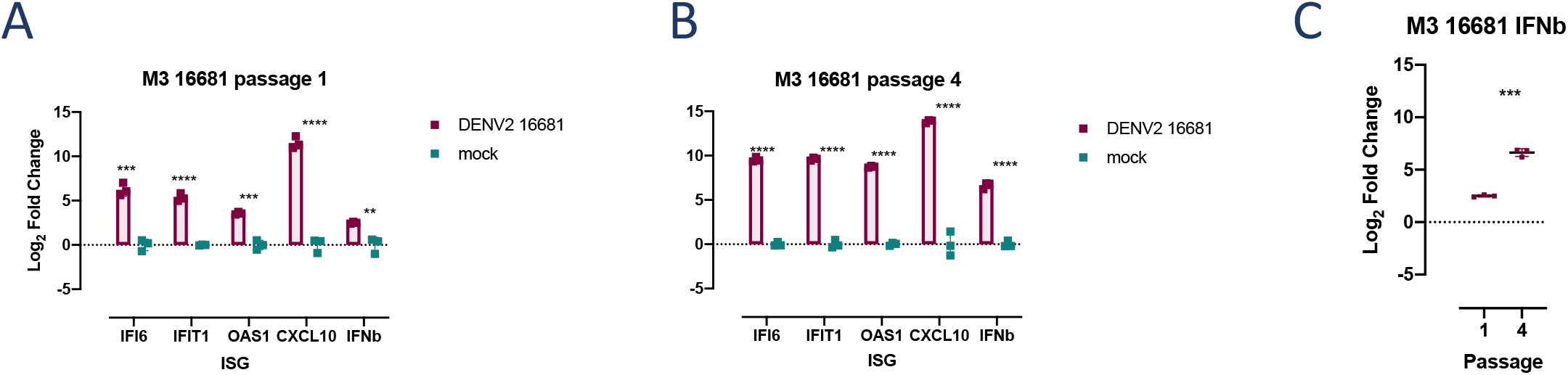
The immune response to DENV2 16681 infection differs with in vitro passage of M3-derived differentiated cells. **(A-B)** Gene expression of IFIT1, IFI6, IFNβ, OAS1, CXCL10, determined by qPCR, during DENV2 16681 infection in P1 (A) and P4 (B) M3-derived differentiated cells with or without DENV2 16681 infection. **(C)** IFNβ expression in infected M3-derived differentiated cells at P1 and P4.

**Supplementary figure 5.**
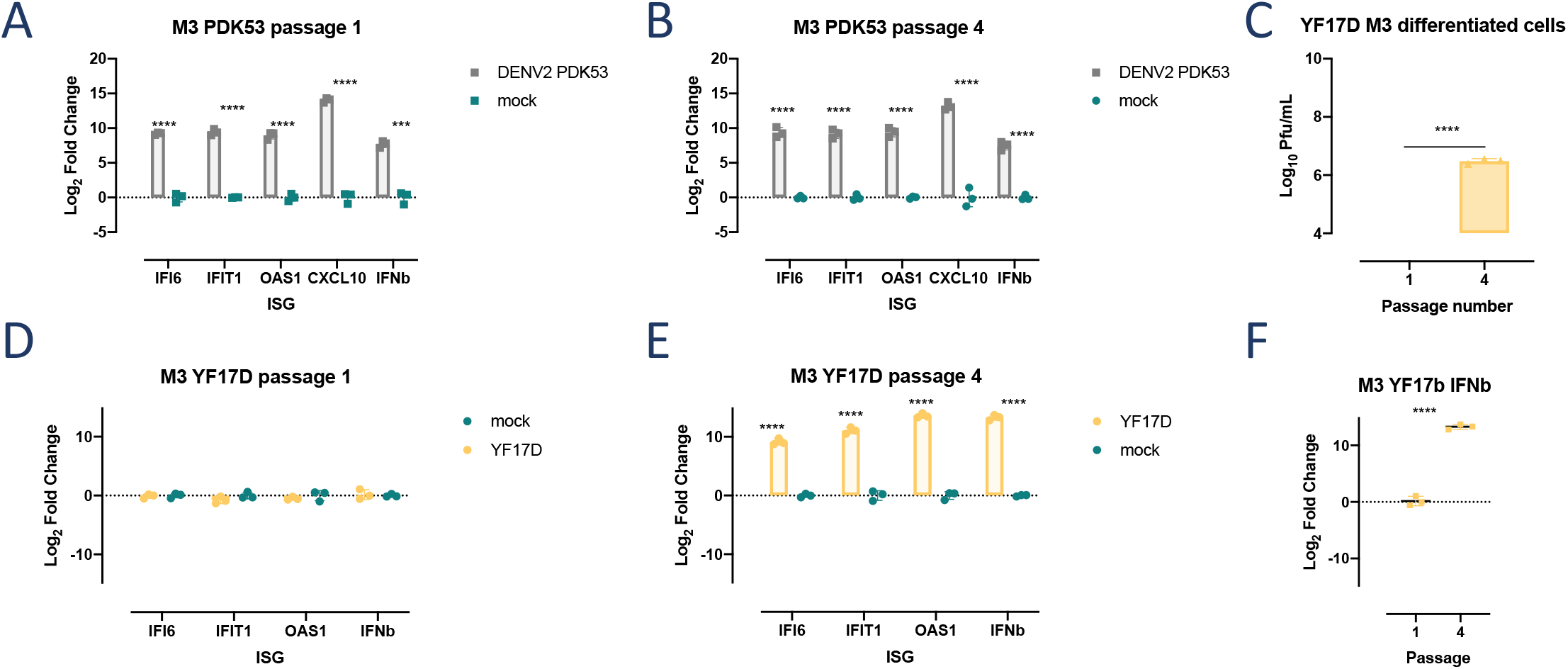
The immune response to DENV2 PDK53 is consistent across passage number while YF17D generates a greater IFNβ at passage 4. **(A-B)** Gene expression of IFIT1, IFI6, IFNβ, OAS1, CXCL10, determined by qPCR following DENV2 PDK53 or mock infection in P1 (A) and P4 (B) M3-derived differentiated cells. **(C)** Plaque titres of YF17D at 48hpi in M3 differentiated cells at passage 1 and 4. **(D-E)** Expression of genes in the canonical IFNβ response pathway (IFIT1, IFI6, IFNβ, OAS1, CXCL10) presented as Log_2_fold change in YF17D infected compared to uninfected cells, at P1 (D) and P4 (D) in M3- and W1-derived differentiated cells. **(F)** IFNβ gene expression at P1 and P4 M3- and W1-derived differentiated cells infected with YF17D and normalized to their respective uninfected controls.

**Supplementary table 1.**
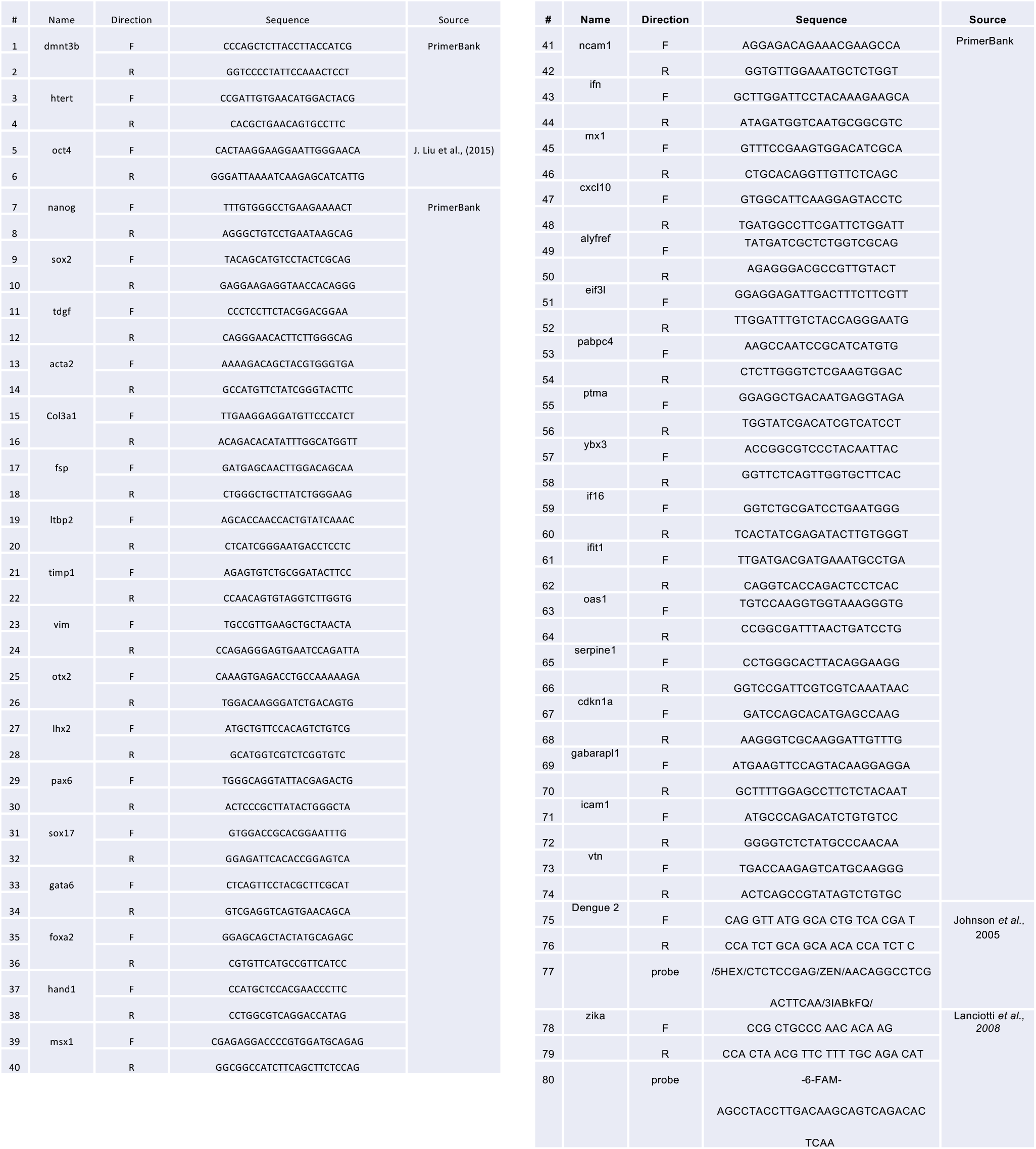
List of qPCR primers.

